# Advancing *Vibrio* genetics: A platform for efficient genomic manipulation

**DOI:** 10.64898/2026.03.10.710856

**Authors:** Jake F. Tatum, Nicholas D. Kraieski, Madeline E. Hamborg, Catherine M. Weatherford, Jackson R. Wells, Carly A. Thatcher, Katherine M. Buckley

**Author notes:** Address correspondence to Katherine M. Buckley.

## Abstract

Most non-model *Vibrio* species lack the genetic tools needed for targeted mutagenesis, which limits the ability to functionally characterize newly identified pathways. To address this challenge, we present here efficient, robust methods for genetically manipulating *Vibrio* species that rely on RecA-mediated homologous recombination and two well-characterized counterselection methods, galactokinase (*galK*) 2-Deoxy-D-galactose (DOG-2) toxicity and the *rpsL*^R^/*rpsL*^S^ streptomycin susceptibility system, both of which are active across a broad range of *Vibrio* species. We further characterized two genus-specific conserved promoters capable of driving high-level ectopic expression across all tested species. These promoters were incorporated into two broadly applicable, conjugatively transferable suicide backbones designed to facilitate double homologous recombination. Using these systems, we successfully disrupted polar flagellar motility in multiple *Vibrio* species and introduced extensive modifications to both flagellar and secretory pathways in *V. diazotrophicus*. Notably, although the *galK* system exhibited broader applicability, the *rpsL* system proved to be more efficient in cases where a streptomycin resistant strain could be generated. We also developed two mobilizable replicative backbones that express pH-stable fluorescent proteins for use within the genus. Collectively, these tools expand the genetic toolkit available for both gene disruption and heterologous gene expression in non-model members of the *Vibrionaceae*.

**Importance:** Members of the *Vibrionaceae* are not only among the most abundant and ecologically influential microorganisms in marine ecosystems, but they also represent major drivers of disease across a wide range of hosts, including humans. However, identifying the genetic determinants of *Vibrionaceae* pathogenesis has remained challenging due to their halophilic growth requirements, restriction enzyme profiles, and resistance to expressing foreign proteins. Canonical counterselection pathways are largely ineffective in these species, which underscores the need for novel and efficient genetic tools to advance functional studies. This study adapts two strategies previously successful in other non-model organisms for use within the *Vibrio* genus and its close relatives. These methods therefore represent an essential step toward overcoming long-standing genetic barriers in *Vibrio* species and provide a framework to expand our understanding of biology in this important bacterial genus.

## Introduction

The genus *Vibrio* (family *Vibrionaceae*: class Gammaproteobacteria: phylum Proteobacteria (1)) includes ∼110 diverse species of heterotrophic bacteria that play important roles in marine ecosystems, including nutrient cycling and nitrogen fixation (2). Although many *Vibrio* species form critical symbioses with plants and animals (3–5), they also serve as important pathogens of both humans and marine organisms. For example, *V. cholerae* is the causative agent of cholera, with over 500,000 confirmed cases and 6,000 deaths globally in 2024 (6). The consumption of seafood contaminated with *V. parahaemolyticus* causes severe gastric distress and *V. vulnificus* is associated with serious skin infections that may progress to necrotizing fasciitis (7–9). Other species such as *V. harveyi* and *V. alginolyticus* are causative agents of many fish-born illnesses and are responsible for substantial economic losses in aquaculture every year (10). In some cases, relationships between *Vibrio* species and hosts are more complex. For example, *V. alginolyticus* is an opportunistic pathogen with significant detrimental effects on aquaculture and global fish populations (11) but serves an important role in preventing disease in the commercially farmed seaweed *Saccharina japonica* (3). The Hawaiian bobtail squid (*Euprymna scolopes*) forms a symbiotic relationship with *Aliivibrio fischeri* within the squid’s light organ (4) whereas larval echinoderms exposed to comparable inoculum doses of the bacteria, exhibit strong immune activation and total mortality (12). The far-reaching physiological, ecological, aquacultural and economic impacts of *Vibrio* species highlight the need to better understand the complex biology of this bacterial genus.

Defining the molecular events that determine the outcome of interactions between *Vibrio* species and hosts is a long-standing challenge due to a lack of efficient genetic tools. Targeted perturbation or elimination of specific genes or gene clusters within bacteria species relies on one of two strategies: 1) introducing exogenous enzymes that manipulate the host DNA directly or 2) taking advantage of endogenous DNA repair enzymes and providing specific “guides” that direct edits based on sequence homology. Methods that use exogenous enzymes rely on transient shuttle vectors that are subsequently cleared from target strains following genomic editing (13, 14). Two of the most common systems encoded on these plasmids are the *E. coli* λ phage-derived Lambda red recombineering cassette and CRISPR-*cas9* (15, 16). Plasmids can be introduced to target strains either by electroporation, which requires electrocompetent target strains, or conjugative transfer, which can be inhibited by a variety of native restriction enzymes or capsule layers (17). These methods often involve complex strategies for identifying rare mutants from background colonies. In contrast, some methods use endogenous enzymes like RecA to drive double homologous recombination via non-replicative suicide backbones. This strategy also requires conjugative transfer or electroporation; however, these backbones are typically smaller and therefore avoid many problems associated with transferring larger exogenous systems into target cells. These methods are best-suited for genetically modifying non-model *Vibrio* species that cannot be efficiently electroporated and are slow to express exogenous proteins (18).

One challenge with relying on endogenous enzymes for genomic editing is that completely eliminating target genes requires two recombination events: the first recombination inserts the backbone into the genome and the second recombination replaces the target gene with a selection cassette. However, these secondary recombination events are rare (19) and therefore require strategies for counterselection against colonies that retain the backbone. Most existing counterselection strategies rely on adding conditionally toxic elements to the backbone. The most widely used of these methods in Gram-negative bacteria utilizes the *Bacillus subtilis* levansucrase gene (*sacB*), which encodes an enzyme that catalyzes the conversion of sucrose into toxic intermediate levans (20). However, this catalytic action is dependent on sucrose concentration and is inactivated by high concentrations of NaCl, rendering it useless in most marine bacteria (including *Vibrio* species) (21). Alternative counterselection strategies that employ toxin-antitoxin pairs have been developed in many bacterial genera (22). However, complications often arise while working with these toxins. In particular, low-level, “leaky” expression during plasmid construction or prior to integration can exert selective pressures that favor inactivating point mutations. Additionally, a lack of a selectable marker to replace target genetic regions increases the amount of satellite colonies that persist from wild type reversions during mutant screening (22). Thus, despite the availability of several counterselection methods to facilitate recombination, their use in *Vibrio* remains challenging.

Here, we present methods for specifically genetically manipulating *Vibrio spp.* that employ two methods of counterselection. First we adapted the galactokinase (*galK*), 2-deoxy-D-galactose (DOG-2) system originally developed for recombineering in *E. coli* (23) for use in *Vibrio spp.* Secondly, we optimized a system that relies on strain-specific streptomycin resistance through point mutations of the *rpsL* gene and its endogenous promoter. The *rpsL* gene and promoter sequences are also broadly conserved throughout the *Vibrio* genus, enabling the use of this counterselection system in any strep^R^ *Vibrio* species. These counterselection systems were introduced into suicide plasmids constructed from the pR6Kg-alpha-1(24) backbone to generate the plasmids pVDOG and pVRPSL. Gene-specific derivates of pVDOG and pVRPSL were used to efficiently eliminate target loci from *V. diazotrophicus*, *V. splendidus*, *V. paucivorans*, *V. coralliilyticus*, and *V. mediterranei*. Additionally, we constructed a suite of replicative plasmids that were stable in *Vibrio* species and can be used for a variety of applications, including fluorescent-labeling (25). Together, our results present a set of genetic tools that are broadly applicable for use in *Vibrio spp.*, do not depend on tightly-controlled induction systems and promote efficient homologous recombination.

## Materials and Methods

### Bacterial strains and culturing methods

Bacterial strains used in this study are summarized in Table 1; complete details can be found in Supplemental Table 1. *Vibrio diazotrophicus* was obtained from ATCC (strain 33466). All *Vibrio* species were grown at 25 °C in Zobell Marine Broth 2216 (MB; BD Difco) and 20 g/L agar as necessary. The conjugative *E. coli* strain RHO3 was a gift from Herbert Schwizer (Addgene bacterial strain #124700) and was cultured at 37 °C in LB supplemented with 2,6-diaminopimelic acid (DAP; 400 µg/mL) to accommodate its Δ*asd* auxotrophy. Unless otherwise stated, antibiotics and supplements were added to growth media as follows: kanamycin (50 µg/mL), chloramphenicol (25 µg/mL), streptomycin (50 µg/mL), tetracycline (10 µg/mL) and/or 0.2% 2-deoxy-D-galactose (DOG-2).

**Table 1:** Summary of strains used in this study.

| Strain | Description | Citation |
| --- | --- | --- |
| ***E. coli*** |  |  |
| EC100D *pir*<sup>+</sup> | *F*<sup>-</sup> *mcrA* $\Delta$(*mrr-hsdRMS-mcrBC*) ϕ80*dlacZ*Δ*M15* Δ*lacX74* *recA1* *endA1* *araD139* $\Delta$(*ara, leu*)7697 *galU* *galK* λ-*rpsL* (*Str*<sup>R</sup>) *nupG* *pir*<sup>+</sup> (*DHFR*) | (78) |
| EC100D *pir*-116 | *F*<sup>-</sup> *mcrA* $\Delta$(*mrr-hsdRMS-mcrBC*) ϕ80*dlacZ*Δ*M15* Δ*lacX74* *recA1* *endA1* *araD139* $\Delta$(*ara, leu*)7697 *galU* *galK* λ-*rpsL* (*Str*<sup>R</sup>) *nupG* *pir*-116(*DHFR*) | (78) |
| RHO3 | Δ*asd thi-1 thr-1 leuB26 tonA21 lacY1 supE44 recA*; integrated RP4-2 *Tcr*::*Mu* Δ*aphA* (λ*pir*<sup>+</sup>) | (79) |
| ***Vibrio diazotrophicus*** |  |  |
| *V. diazotrophicus* | ATCC 33466 | (5) |
| VD::Strep | Derivative of ATCC 33466; Strep<sup>R</sup> due to K42N mutation in *rpsL* | This study |
| VD::*motAB*::*catR* | Replacement of *motAB* genes with *catR* | This study |
| VD::Δ*motAB*::FRT | In frame deletion of *motAB* genes | This study |
| VD::Δ*pomAB*::*catR* | Replacement of *pomAB* genes with *catR* | This study |
| VΔ::*pomAB*::FRT | In frame deletion of *pomAB* genes | This study |
| VD::*flaCD*::*catR* | Replacement of *flaCD* genes with *catR* | This study |
| VD::Δ*flaCD*::FRT | In frame deletion of *flaCD* genes | This study |
| VD::*flaFDAGI/filDS*::*catR* | Replacement of *flaFDAGI/filDS* gene with *catR* | This study |
| VD::Δ*flaFDAGI/filDS*::FRT | In frame deletion of *flaFDAGI/filDS* | This study |
| VD::Δ*hcp*::*catR* | Replacement of *hcp* gene with *catR* | This study |
| VD::Δ*hcp*::FRT | In frame deletion of the *hcp* gene | This study |
| VD::Δ*flaCD*::FRT::Δ*flaFDAGI/filDS*::*catR* | Replacement of the *flaFDAGI/filDS* genes with *catR* in VD::Δ*flaCD*::FRT | This study |
| VD::Δ*flaCD*::Δ*flaFDAGI/filDS*::2FRT | In frame deletion of *flaCD* genes and *flaFDAGI/filDS* genes | This study |
| VD::Δ*pomAB*::FRT::Δ*motAB*::*catR* | Replacement of *motAB* genes with *catR* in VD::Δ*pomAB*::FRT | This study |
| VD::Δ*pomAB*::Δ*motAB*::2FRT | In frame deletion of the *motAB* genes and *pomAB* genes | This study |
| **Other strains used for genetic manipulations** |  |  |
| *V. mediterranei* | Isolated from the mucus of a corallii | JBSQCW01 |
| *V. mediterranei*::Δ*pomAB*::*catR* | Replacement of *pomAB* genes with *catR* | This study |
| *V. coralliilyticus* | Isolated from the mucus of a corallii | JBSXGV01 |
| *V. coralliilyticus*::$\Delta pomAB$::catR | Replacement of *pomAB* genes with catR | This study |
| *V. paucivorans* | Isolated from the mucus of a corallii | JBSXGW01 |
| *V. paucivorans*::$\Delta pomAB$::catR | Replacement of *pomAB* genes with catR | This study |
| *V. splendidus* | ATCC 33125 | (80) |
| *V. splendidus*::$\Delta pomAB$::catR | Replacement of *pomAB* genes with catR | This study |

### Bacterial motility assays

To assess bacterial motility, marine broth plates were prepared as soft agar (0.4% agar) (26). Strains were cultured overnight in MB at 25 °C. Cultures were diluted to OD_600_ = 0.4 and 2 µL were spotted onto the surface of the soft agar MB plates. After 24-96 hours of incubation (25 °C), the diameter of the resultant growth was measured. For consistency, measurements were taken when the wild-type sample reached ∼20 mm in diameter.

### Generating streptomycin-resistant Vibrio strains

Streptomycin-resistant *Vibrio* species were generated as described previously (27). In brief, wild-type species were cultured overnight and spread onto marine broth agar plates (20% agar). Circular disks (6 – 8 mm) of Whatman paper were sterilized and loaded with streptomycin (10 µL of a 50 mg/mL stock solution). A single disk was placed in the center of an inoculated plate. Plates were incubated at 25 °C for 24-36 hours until visible growth appeared surrounding a zone of inhibition caused by the disk. The diameter of the zone of inhibition was recorded (Supplemental Figure 1a). Colonies were collected from the edge of the zone of inhibition and cultured overnight in MB supplemented with sub-inhibitory levels of streptomycin (20 µg/mL). This process was repeated until zones of inhibition were no longer visible around the antibiotic disk.

To confirm that the streptomycin resistance was conferred by mutations in the *rpsL* gene, individual colonies were isolated and cultured overnight at 25 °C in MB supplemented with streptomycin (50 µg/mL). Genomic DNA was extracted and the complete coding sequence of the *rpsL* gene was amplified (primer sequences in Supplemental Table 2). Amplicons were sequenced using Sanger chemistry by Plasmidsaurs (San Francisco, CA).

### Plasmid construction

A summary of the plasmids used in this study is shown in Table 2; complete details, including sequences and Addgene deposition information, can be found in Supplemental Table 3. All primers used for PCR are listed in Supplemental Table 2. PCR reactions were performed using 2X GoTaq Green Master mix (Promega). Plasmids were isolated using ZymoPURE Plasmid kits (Zymo Research). Genomic DNA was isolated using the Wizard Genomic DNA isolation kit (Promega) according to manufacturer’s instructions. Custom DNA fragments were synthesized by Twist Biosciences (San Francisco, CA), cloned into the TOPO TA pCR-4 vector (Invitrogen) and propagated in *E. coli* DH5α. All coding sequences were codon-optimized for expression in *V. cholerae* using the online codon optimization tool from Integrated DNA Technologies (IDT). All ligations were performed overnight at 16 °C using T4 DNA ligase (Invitrogen).

**Table 2:** Abbreviated list of plasmids used in this study.

| Plasmid name | Description | Citations |
| --- | --- | --- |
| **Source plasmids** |  |  |
| pR6Kg-alpha-1 | Original backbone plasmid | (24), Addgene |
| pBAM1 | Source of the oriT sequence | (82) |
| pBACe3.6 | Source of the chloramphenicol resistance gene | (29) |
| pES213 | Replicative plasmid isolated from *Aliivibrio fischeri* | (25) |
| pCMT-*flp* | Flippase shuttle vector | (14) |
| **Integrative plasmids** |  |  |
| pSR6K-1 | Derivative of pR6Kg-alpha-1 with an extended multiple cloning site | This study |
| pSR6KO-1 | Derivative of pSR6K-1 with oriT from pBAM1 | This study |
| pSR6KO-2 | Derivative of pSR6KO-1, KanR replaced with CAT | This study |
| pVF2 | Derivative of pSR6KO-2, with *gfp*<sup>1</sup>, *lac* promoter and RBS | This study |
| pVF3 | Derivative of pVF2 with gene encoding DsRedT.3 DNT gene<sup>1</sup> replacing *gfp* | This study |
| pVFI-1 | Derivative pVF2 with a 500 nt non-coding sequence from of *V. diazotrophicus* chr 2 | This study |
| pVFI-2 – 9 | Derivatives of pVFI with a variety of *V. diazotrophicus* promoter sequences<sup>2</sup> | This study |
| pVKO1 | Derivative of pSR6KO-1 with a single FRT site | This study |
| pVKO2 | Derivative of pVKO1 with an additional FRT site | This study |
| pVDOG | Derivative of pVKO2 with *E. coli galK* and *V. diazotrophicus rpoD* promoter | This study |
| pVRPSL | Derivative of pVKO2 with *V. diazotrophicus* strep<sup>S</sup> variant of *rpsL* and native promoter | This study |
| **Replicative Plasmids** |  |  |
| pVFLR-1 | Derivative of pVF2 with low copy origin | This study |
| pVFHR-1 | Derivative of pVF2 with high copy origin | This study |
| pVFLR-2 | Derivative of pVFLR-1 with the *lac* promoter replaced with *V. diazotrophicus rpoD* promoter | This study |
| pVFHR-2 | Derivative of pVFHR-1 with the *lac* promoter replaced with *V. diazotrophicus rpoD* promoter | This study |
| pVFLR-3 | Derivative of pVFLR-2 with *gfp* replaced with DsRedT.3 DNT<sup>1</sup> | This study |
| pVFHR-3 | Derivative of pVFHR-2 with *gfp* replaced with DsRedT.3 DNT<sup>1</sup> | This study |
| pVFHR-4 | Derivative of pVFHR-2 with *gfp* replaced with *mTurquoise*<sup>1</sup> | This study |
| pVFHR-5 | Derivative of pVFHR-3 with *gfp* replaced with *mScarlet*<sup>1</sup> | This study |
| **Flippase Shuttle Vector** |  |  |
| pREO-1 | Derivative of pSR6KO-1 with the pSC101 origin and the temperature-labile *repA101* | This study |
| pTREO | Derivative of pREO-1 with *tetA* | This study |
| pVCM-*flp* | Derivative of pTREO with *flp* gene<sup>1</sup> and *V. diazotrophicus rpoD* promoter | This study |

<sup>1</sup> Coding sequence was codon-optimized for expression in *Vibrio* strains.
<sup>2</sup> For complete details, see Supplemental Table 3.

### Constructing the pVFI integrative and replicative vectors

pR6Kg-Alpha1 was a gift from Koen Venken (Addgene plasmid #165862) and serves as the backbone for the integrative, suicide and replicative vectors described here. An expanded multiple cloning site (MCS) was introduced to the pR6Kg-Alpha1 using the complementary oligonucleotides pSMCS-F and pSMCS-R (Supplemental Table 2) as previously described (28). In brief, oligonucleotides were resuspended at a final concentration of 80 ng/µL in annealing buffer (10 mM Tris, pH 7.5-8.0, 50 mM NaCl, 1 mM EDTA), mixed in equimolar ratios, heated to 95 °C and allowed to slowly cool to room temperature. The annealed dsDNA fragments were phosphorylated with polynucleotide kinase 1 (pNK1) and ligated into pR6Kg-Alpha1 digested with *SacI* and *SphI* to generate pSR6K-1 (Table 2). To facilitate conjugative transfer, the RP4 origin of transfer (*oriT*_RP4_) was amplified from pBAM1 (pBAM1 was a gift from Victor de Lorenzo; Addgene plasmid #60487) using primers pBAM-*oriT*-F and pBAM-*oriT*-R (Supplemental Table 2). Amplicons were digested and inserted into the *NotI* site of pSR6K-1 to generate pSR6KO-1 (Table 2). A derivative of the pSR6KO-1 backbone was generated using primers pSR6K-*SpeI*-R and pSR6K-*BamHI*-R (Supplemental Table 2) to amplify the MCS and origin sequences excluding the kanamycin resistance gene. A chloramphenicol resistance gene was amplified from pBACe3.6 (29) using primers pCAT-*SpeI*-F and pCAT-*BamHI*-R (Supplemental Table 2). These two fragments were then ligated to generate pSR6KO-2.

Replicative plasmids were generated from pSR6KO-2. A DNA fragment was synthesized consisting of the canonical *lac* promoter (derived from *E. coli*), ribosomal binding site (RBS), GFP coding sequence and restriction enzyme recognition sites. The sequence was inserted between the *XhoI* and *BsrGI* sites in the expanded MCS to generate pVF2. To enable stable replication in *Vibrio* species, DNA fragments were synthesized consisting of the high- and low-copy origin of replication variants of pES213 (25) (Twist Biosciences). Fragments were digested and inserted into pVF2 between the *SalI* and *SacI* sites to generate pVFLR-1 (low-copy replication) and pVFHR-1 (high-copy replication) (Table 2).

The integrative plasmid pVFI-1 was generated from pVF2. The *V. diazotrophicus* genome was analyzed to identify a region that lacked known coding or regulatory sequences (*V. diazotrophicus* strain ATCC 33466 chromosome 2 position: 907618-908119). This target region was amplified using pVDNC-F and pVDNC-R (Supplemental Table 2) and inserted into pVF2 between the *BsrGI* and *AscI* sites. Promoter sequences were identified within the *V. diazotrophicus* genome corresponding to the following genes: *arcA*, *recA*, *motA*, *tusB/C*, *ftsZ*, *rpoD*, *gyrA*, *gyrB*. Promoters were amplified (pVFI2-8-F and pVFI2-8-R; Supplemental Table 2) from genomic DNA and inserted into pVFI-1 in place of the *lac* promoter to generate pVFI-2-9 (Table 1).

The lac promoter in pVFLR-1 and pVFHR-1 was replaced with the *V. diazotrophicus rpoD* promoter to generate pVFLR-2 and pVFHR-2. To allow for differential tagging another DNA fragment was synthesized encoding the dsRedT.3 DNT variant of RFP codon optimized for *V. cholera* (Twist Biosciences) (25). The plasmids pVFLR-3 and pVFHR-3 were generated by replacing the *gfp* sequence in pVFLR-2 and pVFHR-2 with the *rfp* sequence. To increase fluorescent signal, the *V. diazotrophicus gyrB* promoter was used to replace the *rpoD* promoter. To allow for increased fluorescence and faster protein maturation times two additional DNA fragments were synthesized encoding the mScarlet-I3 variant of RFP (30) and the mTurquoise2 variant of GFP (31). Plasmids pVFHR-4 and pVFHR-5 were generated by replacing the gfp and dsRedT.3 DNT coding sequences with of pVFHR-2 and pVFHR-3 with mTurquoise2 and mScarlet-I3, respectively.

### Constructing suicide knockout vectors pVDOG and pVRPSL

The pSR6KO-1 plasmid serves as the backbone for the suicide knockout constructs described here. To add chloramphenicol resistance, the selectable marker used during genomic integration and counter-selection, a DNA fragment was synthesized consisting of the *cat* sequence from pBACe3.6 with a single FRT site upstream of the coding sequence (Twist Biosciences)(29). This fragment was ligated between the *SbfI* and *BsrGI* sites of pSR6KO-1 to generate pVKO1. A second flanking FRT site was added to the 3′ end of the *cat* coding sequence using oligos pFRT-F and pFRT-R (Supplemental Table 2) between the *BsrGI* and *AscI* sites as described above to generate pVKO2 (Table 2).

Two plasmid backbones were generated to facilitate counterselection: pVDOG and pVRPSL. A DNA fragment was synthesized consisting of the minimal promoter of the *V. diazotrophicus rpoD* gene and the *E. coli* galactokinase (*galK*) (Twist Biosciences). This fragment was ligated into pVKO2 to generate pVDOG. pVRPSL was constructed by replacing the *rpoD-galK* cassette of pVDOG with the wild type *V. diazotrophicus rpsL* gene and its native promoter amplified from genomic DNA using primers pVD*rpsL*-F and pVD*rpsL*-R (Supplemental Table 2). For targeted deletions, homology arms were amplified from genomic DNA isolated from the strain of interest using the primers shown in Supplemental Table 2.

Amplified arms were digested and ligated into the *SphI*-*XhoI* and *AscI*-*SalI* restriction sites of either pVDOG or pVRPSL.

### Constructing the flippase shuttle vector pVCM-flp

The previously developed flippase shuttle vector pCMT-*flp*(14) was modified to function more efficiently in *Vibrio* species using the pSR6KO-1 backbone. The *oriV*_R6Kγ_ in this plasmid ensures that the shuttle vector can be replicated in strains of *E. coli* that carry the *pir* gene (*e.g.*, RHO3; Table 2). The pSC101 ori and its temperature-sensitive replication protein *repA101* were amplified from pCMT-*flp* using primers pREP101-F and pREP101-R (Supplemental Table 2) and cloned into pSR6KO-1 to generate pREO-1 (Table 1). The tetracycline resistance gene from pCMT-*flp* was amplified using pTET-F and pTET-R primers and cloned into pREO-1 to generate pTREO. A DNA fragment encoding a N-terminal SUMO- and HIS-tagged flippase (*flp*) and *V. diazotrophicus rpoD* promoter was synthesized (Twist Bioscience) and ligated into pTREO to generate pVCM-*flp* (Table 2).

### Bacterial conjugal mating using the E. coli RHO3 strain

Conjugations were carried out with RHO3 as previously described (32). In brief, RHO3 harboring the mobilizable suicide or replicative backbones were grown overnight at 37 °C in LB broth supplemented with DAP (400 µg/mL) and appropriate antibiotics for plasmid retention.

*Vibrio* strains were grown overnight at 25 °C in marine broth with streptomycin (50 µg/mL) as needed. Cells from 1 mL of both the *Vibrio* and RHO3 overnight cultures were pelleted at 5,000 x g and gently resuspended in 500 µL buffer (10 mM MgSO_4_ for RHO3; 0.2 µm filter-sterile ASW (33) for *Vibrio* strains). Cells were pelleted as above, washed three times in the appropriate buffer and resuspended. Washed RHO3 and the target *Vibrio* strain were mixed and pelleted at a 1:1 ratio to facilitate conjugation. Higher ratios of RHO3:target cells were used (2:1, 5:1 or 10:1) for strains that had poor conjugation efficiency. The resulting pellet was resuspended in 50 µL of ASW and either spotted onto a marine broth agar mating plate supplemented with 400 µg/mL DAP directly or spotted onto a 0.45 µm Whatman filter disk that was placed onto the mating plate. Mating plates were incubated overnight at 25 °C. For mating pairs that were directly plated, a loopful of the colony that arose was streaked for isolation on marine broth agar plates that lacked DAP and were supplemented with chloramphenicol (25 µg/mL) to counter-select against RHO3 and *Vibrio* cells that did not receive the plasmid. Alternatively, filter disks were removed from the mating plates, placed into 1 mL of sterile ASW and vortexed for two minutes. The resulting resuspension was pelleted at 5,000 x g and resuspended in 50 µL of ASW and spread onto chloramphenicol marine broth plates without DAP to isolate transconjugants.

### Counterselection using pVDOG and pVRPSL

Gene-specific pVDOG or pVRPSL plasmids harboring upstream and downstream homology arms were transferred to the target *Vibrio* species using RHO3-mediated conjugation. Genomic integration was verified using an external genome flanking primer and an internal plasmid primer. After successful integration, colonies were passed overnight in marine broth without selection to allow the second recombination event. Serial dilutions were streaked onto MB agar plates supplemented with chloramphenicol (25 µg/mL) and streptomycin (50 µg/mL) for pVRPSL or chloramphenicol (25 µg/mL) and 0.2% DOG-2 for pVDOG. This facilitated counterselection against wild type revertant cells and cells that retained the integrated state.

After counterselection, mutants were verified using PCR and confirmed using whole genome sequencing (Plasmidsaurus) when needed. Genome sequences were compared to wild type using MUMMER (34).

### Sample preparation for scanning electron microscopy

Samples were prepared for scanning electron microscopy (SEM) as described previously (35, 36). In brief, glass coverslips were scored, broken into ∼12 mm diameter pieces and placed into 24-well plates. Bacterial samples were collected in log phase; 100 µL was placed onto coverslips for 30 minutes at room temperature to allow bacteria to adhere. Bacteria were fixed in two steps: 1) 20 µL of 25% glutaraldehyde (Electron Microscopy Sciences, Hatfield, PA, USA) was added slowly in a circular motion to facilitate even mixing and incubated for 30 minutes; 2) 2 mL of 5% glutaraldehyde was gently added to each well and incubated for an additional 30 minutes. Coverslips were washed twice with 0.1% imidazole buffer (pH = 7.0) for 30 minutes.

Cells were first dehydrated by rinsing with increasing concentrations of ethanol (once with 20% ethanol and twice with 30, 50, 75, 80, 90, and 100% ethanol for 30 minutes each). Samples were paused and left overnight in 75% ethanol; subsequent washes were continued the next day.

Samples were further dehydrated using increasing concentrations of hexamethyldisilazane (HMDS; Electron Microscopy Sciences, Hatfield, PA, USA) in ethanol (twice with 20, 50, 70 and 100% HMDS for 30 minutes each). Following the final dehydration step, most of the HMDS was removed from each well and samples were left to airdry uncovered overnight in a fume hood. Coverslips were mounted on stubs and gold-coated for 1 minute on a Quorum Q150R ES sputter coater (Electron Microscopy Sciences). Samples were viewed with a Zeiss EVO50 scanning electron microscope with an accelerating voltage of 20 kV in high vacuum mode. Magnifications up to 50,000X were obtained.

### Phylogenetic analysis

Reference-quality genome sequences were obtained from NCBI: 101 *Vibrio* strains and 8 additional species that served as an outgroup (Supplemental Table 4). Three additional *Vibrio* strains were isolated from mucus samples collected from the Florida false coral (*Ricordea florida*). Using whole-genome sequencing (Plasmidsaurus), these strains were identified as *V. mediterranei, V. coralliilyticus and V. paucivorans*. These genome sequences have been deposited in NCBI (BioProject PRJNA1366596).

Orthologous genes were identified using Orthofinder (37). Two data sets were used to generate phylogenetic trees. First, a tree was constructed using the amino acid sequences of eight genes that were previously identified as single copy orthologs in the *Vibrio* genus: *ftsZ, gyrB, pyrH, dnaJ, rpoA, tyrB, murC, and dnaA* (38) (Supplemental Figure 2). As an alternative, genes present as single-copy orthologs among the species analyzed that encoded the 25 longest protein sequences were selected for analysis (Supplemental Table 4). In each case, amino acid sequences encoded by each gene were aligned using ClustalW (39). Alignments were concatenated and a single maximum-likelihood tree was generated using IQ-TREE 2 (Version 2.1.3)(40). Bootstrap values were calculated based on 1000 replicates modeled using Q. yeast+I+G4. Trees were visualized using phylo.io (41) and iTOL (42).

## Results

### Integrative plasmids can be used to introduce exogenous sequences to Vibrio genomes

As a first step toward designing a system to genetically modify *Vibrio* strains, we constructed the pVFI series of plasmids (Figure 1) to observe expression from integrated backbones within our non-model species. The suicide backbones pSR6KO-1 and pSR6KO-2 can be conjugatively transferred to recipient cells via an origin of transfer (*ori*T_RP4_) (43) but cannot be maintained as extrachromosomal entities in the absence of *pir* expression restricting their host range to specific strains of *E. coli* (*e.g.,* RHO3; Table 1) (44). These plasmids are maintained under either kanamycin or chloramphenicol resistance respectively for flexibility in selection strategies across the genus. Using these mobilizable suicide plasmids originally derived from pR6Kg-Alpha1 (24) (Table 2), we constructed pVFI-1 (Figure 1a) (45, 46). This backbone contains a GFP coding sequence initially expressed from the *E. coli lac* promoter, and a genome targeting sequence (chr2: 907618-908119; CP151843.1; Figure 1b). In the absence of *pir* expression, this targeting sequence promotes Rec-A-mediated recombination to incorporate the introduced DNA into recipient genomes. This process relies on sequence similarity to direct integration (47, 48).

**Figure 1.**
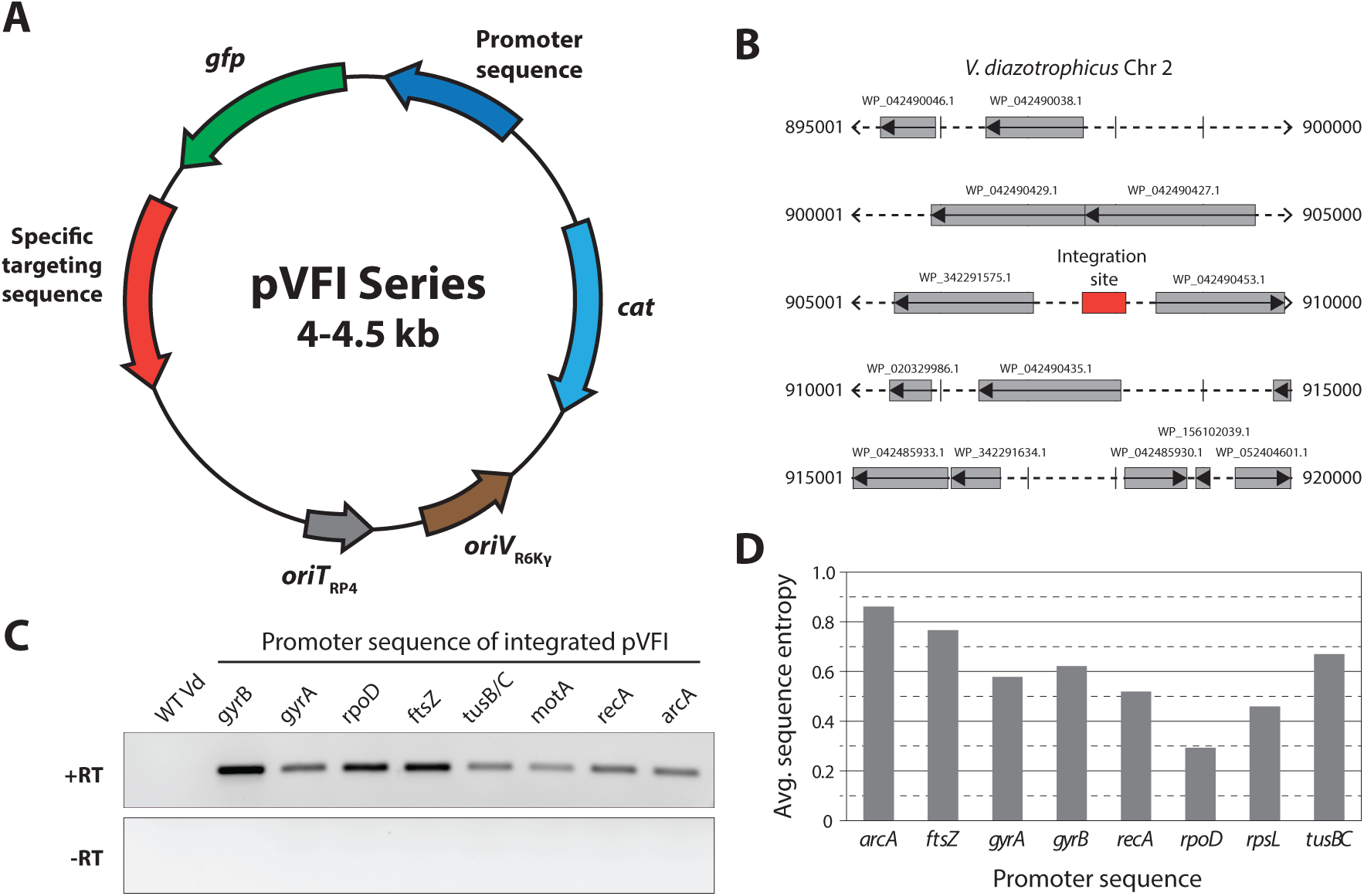
pVFI plasmids enable targeted integration into the *V. diazotrophicus* genome and facilitate exogenous protein expression. A. pVFI plasmids are suitable for conjugative transfer and can be customized with unique targeting, promoter, and coding sequences. A schematic map of pVFI is shown. pVFI plasmids contain the following elements that are flanked by restriction enzyme sites to facilitate modification: 1) a sequence encoding GFP; 2) a promoter sequence, such as the *E. coli lac* promoter; 3) *oriV*_R6Kγ_, a conditionally replicative R6K origin that requires *pir* expression for maintenance; 4) an origin of transfer (*ori*T_RP4_) to facilitate conjugative transfer; 4) a targeting sequence that directs plasmid integration; and 5) the chloramphenicol resistance gene (*cat*). Total plasmid size varies based on the promoter and coding sequences. **B. The integration site does not disrupt known coding or regulatory sequences.** The genomic context of the pVFI integration sequences in *V. diazotrophicus* is shown (red). Neighboring gene sequences are indicated in gray and labeled with NCBI accession numbers. C. Promoter sequences of endogenous *V. diazotrophicus* genes drive variable levels of transcript expression. pVFI plasmids were generated in which the *E. coli lac* promoter was replaced with predicted *V. diazotrophicus* promoter sequences for the following genes: *arcA, ftsZ, gyrA, gyrB, recA, rpoD, rpsL,* and *tusBC*. The expression of *gfp* transcripts was measured using RT-PCR. To ensure that amplification was not due to genomic DNA contamination, samples lacking reverse transcriptase (-RT) are shown. **D. Predicted promoter sequences exhibit differential levels of sequence conservation.** Sequences upstream of the start codon (500 nt or the distance to a neighboring gene) were isolated from 20 representative *Vibrio spp.* (Supplemental Table 4). Sequence diversity was calculated as Shannon’s entropy for each region such that higher values correspond to increased sequence variation.

In this aim, expression from the lac promoter in pVFI-1 a pSR6KO-2 derivative was tested as a candidate for ectopic expression of counter-selective markers from integrative backbones. Following RHO3 mediated conjugal mating to *V. diazotrophicus,* integrants were confirmed using genomic PCR and whole genome sequencing to initially generate VD::pVFI-1. The genomic insertion site was selected by identifying a 500 nt region that lacks known coding sequences or regulatory elements (chr2: 907618-908119; CP151843.1; Figure 1b). Comparison of the growth curves VD::pVFI-1 vs. wild-type *V. diazotrophicus* indicate that using this integration site did not disrupt normal cell growth (Supplemental Figure 3). Although RT-PCR revealed *gfp* transcripts were expressed at high levels in VD:pVFI-1, we were unable to detect substantial signal via fluorescent microscopy (data not shown). We hypothesized that this may be due to insufficient expression levels driven by the *E. coli lac* promoter and therefore investigated a suite of native promoters from *V. diazotrophicus* for conservation and use across the genus.

### The conserved rpoD promoter drives constitutive expression in Vibrio species

To support counter-selection strategies, we identified conserved promoter sequences that can drive differential expression in *V. diazotrophicus* and other *Vibrio* species. Predicted regulatory regions were isolated from eight operons encoding genes involved in normal cellular physiology: *arcA, ftsZ, gyrA, gyrB, motA, rpoD, recA,* and *tusB/C*. Sequences were amplified, inserted into pVFI in place of the *lac* promoter (generating pVFI2-9; Table 2) and integrated in the *V. diazotrophicus* genome (VD::pVFI2-9; Supplemental Table 1). RT-PCR revealed that each of the promoter sequences induced expression of *gfp* transcripts at varying levels in the integrated strains (Figure 1c). Because we were interested in identifying promoters with broad activity within the genus, we analyzed the sequence conservation of the promoters for each of the genes in 20 representative *Vibrio* species (Supplemental Table 4). The most conserved of these promoters is that of *rpoD* (Figure 1d). The predicted ribosomal binding site (RBS) and transcriptional binding sites (-35 and -10 regions) were also highly conserved, exhibiting almost no variation among the *Vibrio* sequences. Together, these findings suggest that the *rpoD* promoter is a highly conserved regulatory element across *Vibrio* species, making it a promising candidate for driving consistent gene expression across this genus.

### Replicative plasmids can be used to stably express genes of interest in Vibrio

As an alternative strategy to express exogenous sequences, we were interested in developing replicative plasmids using these characterized promoter sequences (Figure 1c) that can be stably maintained in *Vibrio* species without chromosomal integration (Figure 2). For this, we used the *gyrB* promoter, which is less conserved than that of *rpoD*, but drove higher expression (Figure 1c,d). A replicative backbone (pVFHR-1; Table 1) was generated from pVF2 by adding the high-copy origin of replication from pES213 that is permissive to *Vibrio* species (*oriV_Vibrio_*) (25). Previous work labeling *Vibrio* species relied on differential staining (49), or the expression of sequences encoding either GFP (25, 50) or DsRed/RFP variants (51, 52). However, we were interested in generating a strain that expresses a fluorescent protein with a short maturation time and is stable in a range of pH conditions. We therefore introduced sequences (codon-optimized for expression in *Vibrio* species) that encode either mTurquoise2 (a GFP-derived cyan fluorescent protein; (53)) or mScarlet-I3 (a synthetic derivative of RFP; (54)) into the pVFHR-derived backbones to generate pVFHR-4 and pVFHR-5 (Table 2; Figure 2a). These replicative plasmids were conjugatively introduced to *V. diazotrophicus*; the resulting strains were passaged over five growth cycles in the presence and absence of selective antibiotics to assess stability and expression. Consistent with previous analysis of plasmids containing the pES213 origin (25, 46), plasmid loss was minimal and expression was maintained (data not shown). RT-qPCR confirmed that, in *V. diazotrophicus* the expression of the fluorescent protein transcripts was 4.6 – 46.8X higher in strains containing replicative plasmids (pVFHR-4 and pVFHR-5) relative to integrated plasmids (Figure 2b). Additionally, strains harboring these replicative plasmids could be easily visualized using fluorescent microscopy (Figure 2c).

**Figure 2.**
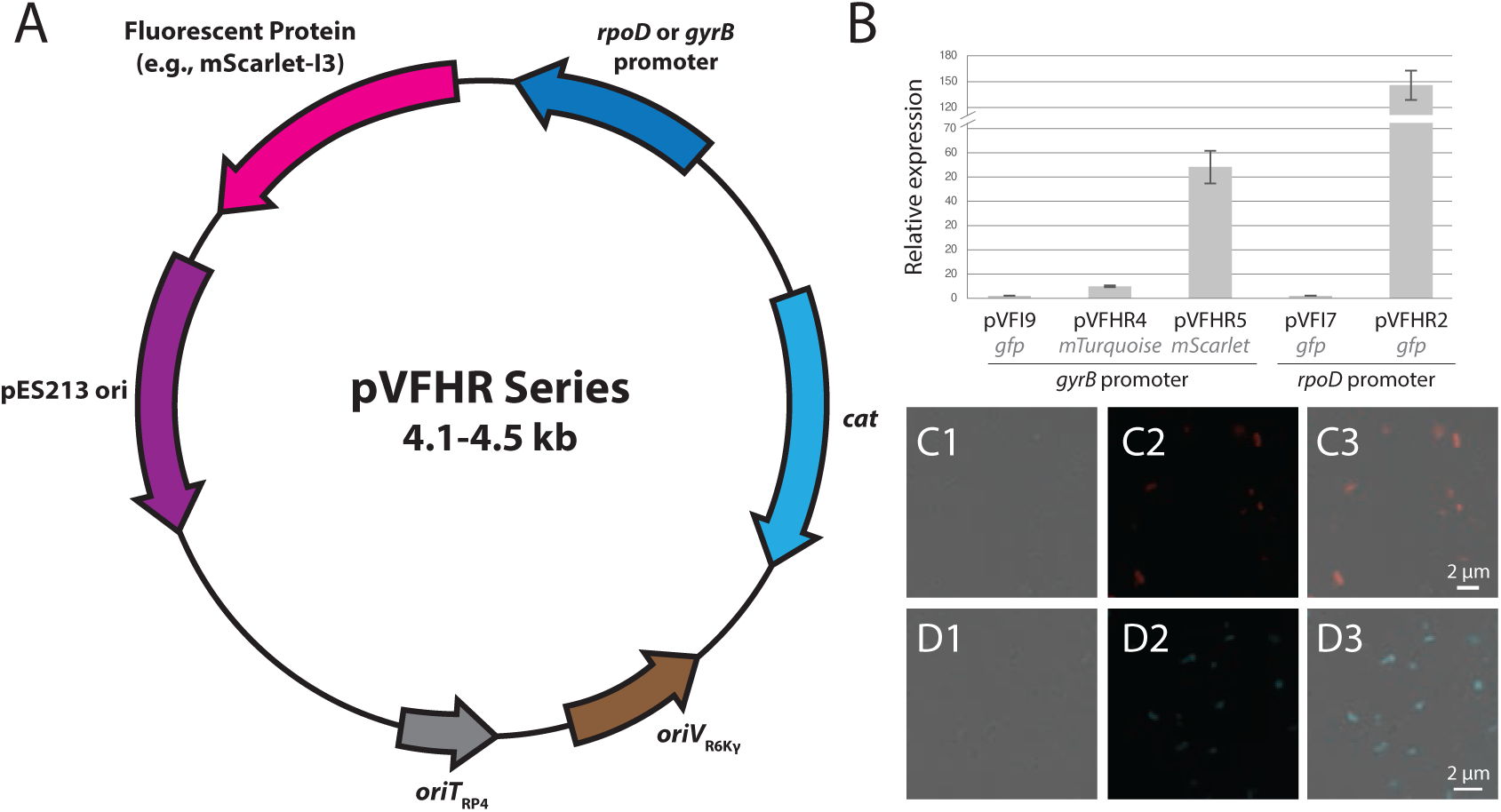
The pVFHR replicative plasmids can be optimized for different expression levels. **A. The vector map of the pVFHR plasmids is shown.** All elements shown in this map are flanked by common restriction enzyme recognition sites and can be altered based on specific needs. Here, we have introduced either the *rpoD* or *gyrB* operon promoter sequences from *V. diazotrpohicus* as well as a sequence encoding the pH-stable fluorescent proteins mScarlet or mTurquoise. B. **Replicative plasmids promote higher levels of transcription than integrative**. qPCR was used to quantify transcript expression in *V. diazotrophicus* strains that either carried pVFHR replicative plasmids or integrated exogenous sequences into the genome using the pVFI plasmids (Figure 1). Results indicate that expression is much high in strains containing the replicative plasmids as previously seen with the pES213 origin. Relative expression levels were calculated using the ΔΔCt method normalized to *pomA* as a reference gene. Reactions were run in triplicate; error bars represent variation among technical triplicates. **C, D. Fluorescent protein is detectable in *V. diazotrophicus* tagged with pVFHR plasmids**. DIC (C1, D1), fluorescent (C2, D2) and overlay (C3, D3) images are shown. Images were captured using a 100X objective; scale bars represent 2 μM.

Together, these results indicate that the pVFHR replicative plasmids can be used to generate strains that stably express exogenous proteins however, these levels of expression were not achievable by integrated backbones.

### Suicide plasmids and counterselection enable targeted chromosomal deletions in Vibrio genomes

Using these broadly recognizable promoters, we can stably express ectopic genes from integrated backbones at a range of levels, providing the flexibility needed to adapt new counterselection strategies across the genus. Implementing these approaches requires two sequential RecA-mediated homologous recombination events to generate clean deletions (Figure 3A)(55). In the first event, the entire plasmid backbone integrates into the chromosome through one of the homology arms. The second recombination event between the remaining arm and the host genome is considerably rarer. As a result, numerous counterselection strategies have been developed to eliminate integrants that retain the full backbone and to enrich for the small population of double-recombinant clones.

**Figure 3.**
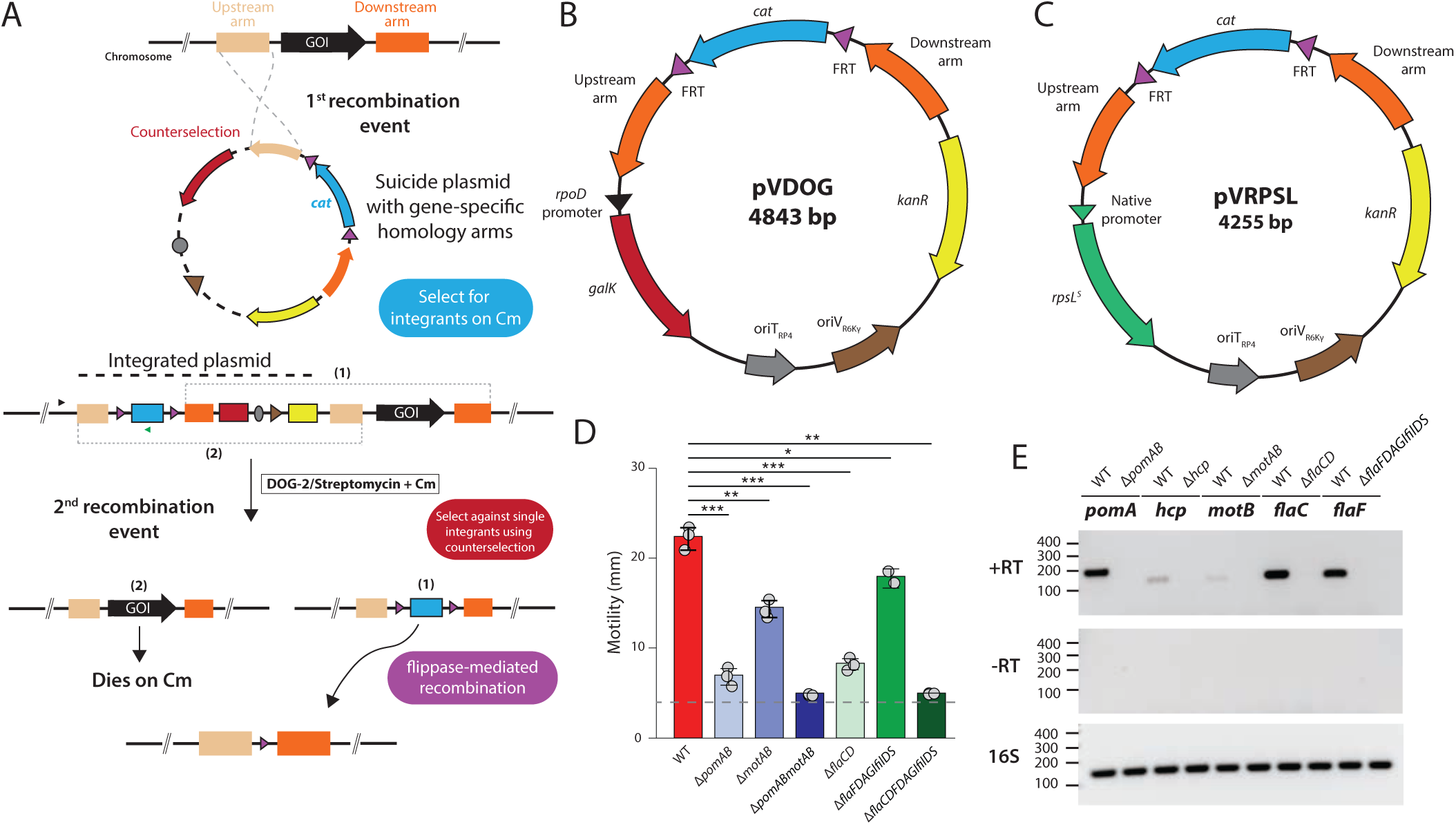
The pVRPSL and pVDOG suicide plasmids provide efficient counterselection and enable targeted gene deletion. A. Rec-A mediated gene deletions require two sequential recombination events. A schematic of the two-step allelic exchange process mediated by pVRPSL or pVDOG is shown. Integration can occur with roughly equal probabilities on either side of the homology arm that directs recombination around the target gene; for simplicity, the upstream arm is illustrated here. Integrants are identified by antibiotic resistance, and integration is confirmed using a combination of integration-specific (p17 F/R) and genomic primers (the upstream flanking F and p17-R primers are depicted). Successful excision and mutant recovery require recombination through the second homology arm. Only strains that have lost the pVRPSL or pVDOG backbone can survive on streptomycin- or DOG-2-supplemented media. Although plasmid loss theoretically yields both wild-type and mutant colonies in equal proportions, inclusion of chloramphenicol selects exclusively for mutant isolates. If desired, the chloramphenicol cassette can subsequently be removed using the pVCM-*flp* shuttle vector (Figure 4). B. The pVDOG vector encodes *galK* to enable DOG-2 mediated counterselection. The vector map is shown. This plasmid contains the sequencing encoding the *E. coli* galK enzyme that phosphorylates DOG-2 to form the metabolically inert 2-deoxy-D-galactose-1-phosphate, which is toxic at high concentrations in Gram-negative bacteria (23, 56). C. The pVRPSL vector encodes streptomycin-susceptible *rpsL* to mediate counterselection. The vector map is shown. D. Motility assays were used to verify the loss of movement-associated genes in *V. diazotrophicus*. The initial inoculum (4 mm) is indicated by a dashed gray line. All assays were performed in biological triplicates; error bars indicate the standard deviation. Statistical significance was calculated compared to wild type strains by Students t-test (***, p < 0.001; **, p < 0.01; * p < 0.05; ns, p > 0.05). **E. RT-PCR confirms loss of transcription in deletion strains.** Expression was quantified in wildtype (WT) and single-locus mutants of *V. diazotrophicus* generated using pVDOG or pVRPSL (Δ*pomAB*, Δ*hcp*, Δ*motAB*, Δ*flaCD*, Δ*flaFDAGIfilDS*; see Supplemental Table 1 for strain information). Primer sequences are listed in Supplemental Table 3.

Identifying the rare clones that have undergone the second recombination requires an efficient, controllable system for counter-selection. Equipped with a suite of integrative promoters capable of specifically controlling gene expression (pVFI-1-9), we investigated the possibility of adapting the well-characterized galactokinase (*galK*) system (23) for efficient counterselection in *Vibrio spp*. GalK phosphorylates α-D-galactose to form galactose 1-phosphate as part of galactose catabolism. This enzyme also phosphorylates the substrate 2-deoxy-D-galactose (DOG-2) to form the metabolically inert 2-deoxy-D-galactose-1-phosphate, which is toxic at high concentrations in Gram-negative bacteria (23, 56). Because GalK has a low affinity for DOG-2, elevated expression of the enzyme is required to permit accumulation of the toxic intermediate (57).

To build an integrative backbone that enables counterselection, we inserted the *E. coli galK* coding sequence and the broadly conserved *rpoD* promoter from *V. diazotrophicus* (Figure *2*e) to produce pVDOG (Figure 3b; Table 2). The *rpoD* promoter was selected to avoid the overexpression-associated stress that can impair counterselection efficiency. The *E. coli* version of *galK* was used rather than a *Vibrio* homolog to maintain substrate specificity (56). The resulting pVDOG backbone includes ∼500 nt upstream and downstream homology arms that flank a chloramphenicol resistance cassette, enabling targeted recombination and replacement of specific genomic loci.

To evaluate this system, we targeted genes involved in motility, which produces a distinct and easily measurable phenotype. In *V. diazotrophicus*, motility is driven by two flagellar systems: 1) a single polar flagellum powered by sodium ion flux; and 2) lateral flagella energized by canonical proton motive force (5). To disrupt polar flagellar function, we targeted *pomA* and *pomB*, which encode stator proteins responsible for generating torque from sodium ion gradients. (Table 1) (58, 59). Plasmids carrying the corresponding homology arms were introduced into *V. diazotrophicus* via conjugation, and transconjugants were isolated. Counterselection was performed by supplementing culture media with 0.2% DOG-2. Successful genomic exchanges in the resulting strain (VD::Δ*pomAB*::*cat*) were verified by PCR and whole-genome sequencing to rule out off-target effects (Figure 3; Table 2). Motility assays confirmed that VD::Δ*pomAB*::cat exhibited significantly reduced motility compared to the wild-type strain (average colony diameters: 1.98 cm vs. 0.86 cm respectively; Figure 3d). These results indicate that the *galK*-based counterselection system reliably eliminates target genes and produces the expected phenotypic changes in *V. diazotrophicus*.

One potential limitation of the *galK* system is that strains may require higher concentrations of DOG-2 to maintain comparable recombination frequency over time (57) and mutations in *galK* may lead to inactivation (56, 57). Although the lower expression levels driven by the *rpoD* promoter should mitigate these issues, it can still complicate the ability to sequentially edit multiple loci within a single strain. To address these challenges, we developed pVRPSL (Figure 3d; Table 2), which relies on integrant-dependent streptomycin susceptibility for counterselection. We first generated a streptomycin-resistant *V. diazotrophicus* strain (VD::*rpsL*^R^; Table1) by serially passaging through sublethal streptomycin concentrations.

Resistance typically arises from mutations in the single-copy gene *rpsL*, which encodes ribosomal protein S12 (60, 61). PCR and Sanger sequencing were used to confirm that VD::*rpsL*^R^ carries a K42N mutation (Supplemental Figure 1b). The pVRPSL plasmid encodes the wild-type, streptomycin-susceptible variant of *V. diazotrophicus rpsL* (*rpsL*^S^) and its native promoter in place of *galK* (Figure 3d). Using the endogenous promoter ensures that, upon integration, both the plasmid-derived *rpsL^S^* and chromosomal *rpsL^R^* alleles are expressed at comparable levels, thereby rendering integrants susceptible to streptomycin.

To assess the efficacy of genome editing with pVRPSL, we targeted the type VI secretion system (T6SS)(5). The T6SS depends on the single-copy *hcp* gene, which encodes the structural subunit that polymerizes to form the molecular needle that is fired into target cells (62). Gene-specific homology arms were cloned into pVRPSL (Table 2), and the resulting plasmid was introduced to VD::*rpsL*^R^ via conjugative transfer. Integrants were identified and counterselection was performed on streptomycin-supplemented media. Successful mutants (VD::Δ*hcp*::*cat*) were verified by genomic PCR and RT-PCR (Figure 3e).

One potential limitation of both the pVDOG and pVRPSL systems is that their backbones contain sequences homologous to endogenous *Vibrio* genes (*i.e.*, the *rpoD* promoter and *rpsL* coding sequence). Although these systems efficiently generate the intended mutants, we occasionally observed low-frequency off-target transconjugants, particularly with pVRPSL (∼1 in 20 colonies tested). Accordingly, all transconjugants should be verified using PCR with gene-specific primers before proceeding to counterselection (Supplemental Table 3; Figure 3a).

### A shuttle vector encoding flippase enables nearly seamless mutagenesis

Analysis of VD::Δ*pomAB*::*cat* revealed that, although motility was reduced compared to wild-type strains, this mutant was not completely immotile (Figure 3c). We hypothesized that this residual movement was driven by the lateral flagellar system. Rotation of these flagella requires the MotA and MotB proteins, which form the proton dependent stator complex. To delete these genes from the VD::Δ*pomAB*::*cat* background using pVRPSL or pVDOG, it was first necessary to remove the chloramphenicol resistance cassette (*cat*) that replaced the *pomAB* locus. For this, the *cat* gene in both pVDOG and pVRPSL is flanked by flippase recognition target (FRT) sites (Figure 3a, b). Flp recombinase binds to these sites and catalyzes excision of the intervening DNA, leaving a single, 34 nt FRT scar (63). This strategy enables removal of selection markers from mutant strains and facilitates subsequent genetic modifications.

To transiently express *flp*, we generated a *Vibrio-*specific conditionally replicative plasmid. This vector, pVCM-*flp,* was derived from pCMT-*flp* (14), and incorporates several features to optimize performance in *Vibrio* species (Figure 4; Table 1). pVCM-*flp* enables efficient passage through *pir^+^ E. coli* strains and carries a *Vibrio* codon-optimized *flp* gene regulated by the conserved *V. diazotrophicus rpoD* promoter (Figure 1e). This promoter was selected to moderate expression, as Flp is a large, yeast-derived protein that could impose a metabolic burden on recipient cells. Additionally, pVCM-*flp* includes a temperature-dependent origin of replication (pSC101 ori; (64)) in which a mutated form of *repA101* is unstable at 37 °C. This feature facilitates plasmid clearance following flippase-mediated recombination.

**Figure 4:**
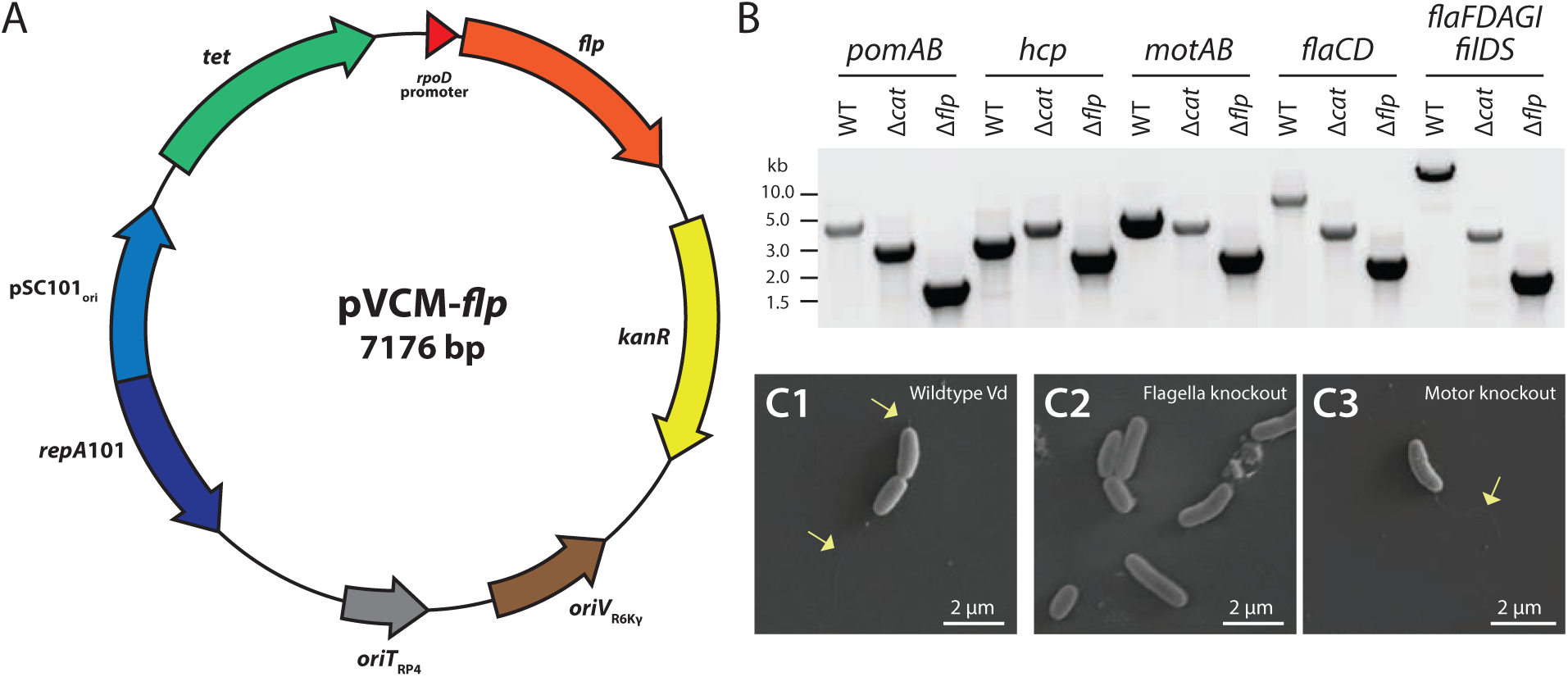
Flippase-mediate recombination can be used to generate nearly scarless genomic sequences. A. pVCM-*flp* is a temperature-sensitive, replicative plasmid that encodes flippase. The vector map is shown. pVCM-*flp* contains a codon-optimized transcript encoding flippase under the control of the *V. diazotrophicus rpoD* promoter. The temperature-sensitive oriR101 origin or replication enables plasmid curing by culturing at 37 C. **B. Verification of flippase-mediated recombination by PCR.** The genomic region surrounding the target genes was amplified in wild type (WT), mutant (Δ*cat*), and flipped strains (Δ*flp*) of *V. diazotrophicus* (Δ*pomAB*, Δ*hcp,* Δ*motAB*, Δ*flaCD*, and Δ*flaCDFDAGIfilDS*). C. *V. diazotrophicus* mutants that lack flagellar motors retain flagella but lose motility. pVCM-*flp* plasmids were used to eliminate counterselection sequences and delete multiple loci in *V. diazotrophicus* (Vd) strains. SEM images are shown for the wildtype, flagella knockout (VD::Δ*flaCD*::Δ*flaFDAGI*/*filDS*::2FRT) and motor knockout (VD::Δ*pomAB*::Δ*motAB*::2FRT) strains. Polar flagella are observed on both wildtype and motor knockout strains (yellow arrows) but absent from flagella knockout *V. diazotrophicus.* Scale bars indicate 2 μM.

pVCM-*flp* was conjugatively transferred to VD::Δ*pomAB*::*cat* to generate the near-seamless mutant VD::Δ*pomAB*::FRT (Table 1). To generate a fully non-motile strain of *V. diazotrophicus,* the *motAB* locus in VD::Δ*pomAB*::FRT was targeted for deletion using a pVRPSL backbone (Table 1). The double mutant strain (VD::Δ*pomAB*::Δ*motAB*::*cat*::FRT) was isolated following counterselection and confirmed by PCR and genome sequencing (Figure 3d). The *cat* gene was subsequently removed by the pVCM-*flp* vector to generate the near-seamless double mutant VD::Δ*pomAB*::Δ*motAB*::2FRT strain (Table 2). Motility assays revealed that this mutant did not move past the initial inoculum (0.4 cm; Figure 3c).

As an alternative approach to eliminating motility in *V. diazotrophicus*, we targeted the genes encoding the flagellin subunits. Flagellin subunit genes are usually present in multiple copies throughout bacterial genomes, and each contribute to the polymerization of the filament once transferred outside of the cell through the flagellar apparatus (65). These loci (*flaCD* and *flaFDAGI/filDS*) were identified within the *V. diazotrophicus* genome and targeted for deletion using gene-specific pVRPSL plasmids (Table 2). Single locus mutant strains were generated (VD::Δ*flaCD*::FRT and VD::Δ*flaFDAGI/filDS*::cat; Table 2), each of which exhibited decreased motility relative to wild type (Figure 3c). We subsequently generated the double mutant strain (VD::Δ*flaCD*::Δ*flaFDAGI/fliDS*::2FRT) using pVCM-*flp*. As with VD::Δ*pomAB*::Δ*motAB*::2FRT, this strain was entirely non-motile (Figure 3c).

To further validate the motility phenotypes, we examined the external flagellar structures of the two non-motile strains (VD::Δ*flaCD*::Δ*flaFDAGI/fliDS*::2FRT and VD::Δ*pomAB*::Δ*motAB*::2FRT) and the wild-type strain using scanning electron microscopy (SEM). In wild-type cells, both lateral and polar flagella were readily observed (Figure 4).

Flagella were also visible in the double stator knockout strain (VD::Δ*pomAB*::Δ*motAB*::2FRT; Figure 4d), indicating that although these cells can assemble flagella, the structures are non-functional. In contrast, the filament knockout strain (VD::Δ*flaCD*::Δ*flaFDAGI/fliDS*::2FRT) lacked any external structures resembling flagella (Figure 4d). No additional morphological differences were observed among the three strains. Together, these findings corroborate the motility assays and demonstrate the precision and efficacy of the recombineering methods described here to manipulate *V. diazotrophicus*.

### The pVDOG/pVRPSL plasmids can be broadly used among Vibrio species

Given the broad conservation of the *rpoD* and *rpsL* promoters among *Vibrio* species (Figure 1e), we investigated the possibility that the pVDOG/pVRPSL backbones could be used to modify genomes in other *Vibrio* strains. To provide a phylogenetic context for selecting candidates, we reconstructed the evolutionary relationships among 101 *Vibrio* strains (Figure 5a). All whole genome sequences that were assembled at scaffold-level higher, were retrieved from NCBI (Supplemental Table 4). From these genomes, the 25 longest single copy orthologs were identified and used to reconstruct the evolutionary history of this genus (Figure 5). The resulting tree was consistent with those generated using only eight genes that are commonly used for phylogenetic analyses (Supplemental Figure 2), providing confidence in the robustness of the selected marker set. To determine if the pVDOG and pVRPSL plasmids can mediate genomic editing across *Vibrio* species, candidate strains were selected that span a broad phylogenetic range: *V. mediterranei, V. coralliilyticus, V. paucivorans,* and *V. splendidus* (Figure 5; Table 1).

**Figure 5.**
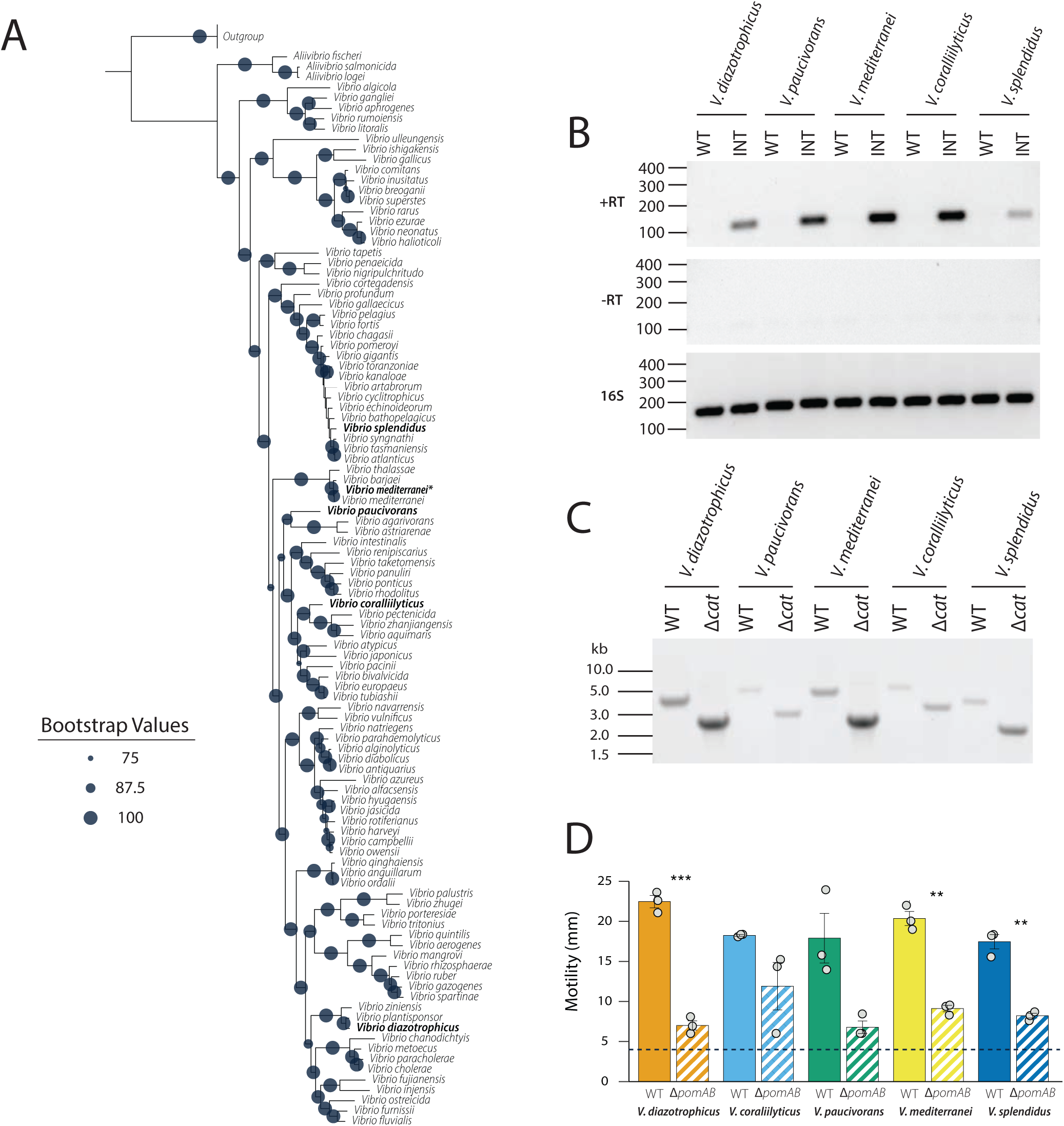
The pVDOG/pRPSL plasmids are effective across a broad range of *Vibrio* species. **A. The phylogenetic relationships among *Vibrio* species were inferred using single-copy orthologs.** A maximum-likelihood tree was constructed using the amino acid sequences of the 25 longest orthologs identified across all *Vibrio* genomes analyzed (Supplemental Table 4). This topology is consistent with that generated using eight commonly used orthologs (Supplemental Figure 2). *Vibrio* species successfully edited using the pVDOG/pVRPSL plasmids are highlighted in bold; the laboratory strain of *V. mediterranei* used is denoted with an asterisk. Outgroup species include *Escherichia coli, Klebsiella pneumoniae, Salmonella enterica, Serratia marcescens,* and *Yersinia pestis.* Bootstrap values shown were calculated using 1,000 replicates; only values greater than 75 are shown. **B. The *rpoD* promoter dives high-level expression from integrated pVDOG backbones.** RT-PCR was performed on *V. diazotrophicus*, *V. paucivorans*, *V. mediterranei*, *V. coralliilyticus*, and *V. splendidus* strains carrying single pVDOG integrations using primers specific to the *E. coli galK* gene. RT loading controls are depicted below each sample. **C. Genomic PCR confirms deletion of *pomAB* across multiple *Vibrio* species.** The chromosomal regions flanking the *pomAB* loci were amplified from wild-type (WT) and mutant (Δ*cat*) strains of *V. diazotrophicus*, *V. paucivorans*, *V. mediterranei*, *V. coralliilyticus*, and *V. splendidus*. **D. Motility assays demonstrate the broad applicability of this system across *Vibrio* strains.** In each species tested, deletion of *pomAB* genes reduced motility. Colony diameters were measured, with the initial inoculum (4 mm) indicated by the dashed line. All assays were performed in biological triplicates; error bars indicate standard deviation. Statistical significance was calculated compared to wild type strains using Students t-test (***, p < 0.001; **, p < 0.01; * p < 0.05; ns, p > 0.05).

For this aim, we targeted the *pomAB* locus in each species, as eliminating these genes results in easily measurable phenotypes in motility assays. pVDOG and pVRPSL backbones were generated with the corresponding homology arms for the *pomAB* loci in each species. Plasmids were introduced to candidate strains via conjugative transfer; integrants were selected and confirmed using PCR and whole genome sequencing. We first assessed the activity of the *V. diazotrophicus rpoD* promoter from the pVDOG backbones in each of the candidate species by quantifying *galK* expression (Figure 5b). To enable pVRPSL mediated counterselection, streptomycin resistant strains were generated for each species (Table 2). Mutations in each of the *rpsL* genes were verified by PCR and sequencing (Supplemental Figure 1c). Using methods described above, Δ*pomAB* strains were generated for each of the five species and confirmed using PCR (Figure 5b). Importantly, motility defects were observed for each of the mutant strains relative to their wildtype counterparts, which is consistent with deletion of the genes encoding the pomAB flagellar stators (Figure 5c). Notably, the pVRPSL backbones had a higher mutant yield than the pVDOG backbones, which may be due to variation in DOG-2 import and spontaneous mutations that may arise in the *galK* coding sequence (66). Together, these results indicate that pVRPSL and pVDOG are broadly applicable for genetically manipulating *Vibrio* species and, possibly, other genera within *Vibrionaceae*.

## Discussion

We describe here an efficient and adaptable suite of tools for the genetic modification of non-model species within the *Vibrionaceae*. The conjugatively transferable plasmids pVDOG and pVRPSL, which incorporate counterselection strategies to enable targeted gene deletion, and pVCM-*flp*, which encodes flippase to allow nearly scarless editing, represent refined applications of existing techniques to facilitate RecA-mediated recombination in diverse *Vibrio* genomes. In addition, the pVFLR and pVFHR replicative plasmids provide a method for rapidly tagging *Vibrio* strains using fluorescent proteins. Together, these tools open new avenues of study into non-model *Vibrio* species.

Genetic manipulation of species within the *Vibrionaceae* can be challenging within clades that are phylogenetically distant from well-studied taxa such as *A. fischeri, V. cholera* or *V. vulnificus*. Several factors contribute to these experimental difficulties: 1) most *Vibrio spp.* are strictly halophilic, which interferes with electroporation efficiency (67, 68); 2) many species secrete extracellular and periplasmic nucleases that degrade incoming DNA and impede uptake (17, 69), and 3) many species within this genus are not naturally competent in the absence of genetic modifications (70). We therefore focused our attempts to genetically modify non-model *Vibrio* species on strategies that rely on conjugative transfer.

*Vibrio* strains also pose challenges during counterselection steps; SacB is inefficient under high-salinity growth conditions, and sucrose-resistance phenotypes are frequently observed among integrants (21). Induced expression of toxins from toxin:antitoxin modules on integrated backbones is widespread in recombineering across bacterial phyla (22, 71–73). However, these coding sequences are frequently subject to inactivating mutations during cloning or within recipient cells which, in our hands, significantly reduced the recovery of the desired mutants. Consequently, we here describe the adaptation of two established counterselection strategies (*rpsL-*mediated streptomycin resistance and galK/DOG-2 toxicity) to the *Vibrio* genus, creating recombineering systems that enable rapid and reliable genetic manipulation in previously recalcitrant or understudied species. We further demonstrate sequential applications of these methods in *V. diazotrophicus* by creating multiple single- and double-gene knockout strains.

Our initial attempts to perform double homologous recombination yielded numerous background colonies representing unsuccessful recombinants that escaped counterselection, even in cases where recombination at one homology arm was confirmed by PCR. We determined that *galK* expression driven by the *lac* promoter was insufficient to support robust DOG-2–based counterselection, even under inducing conditions. To overcome this limitation, we identified conserved promoter elements from *V. diazotrophicus* (*i.e., rpoD* and *gyrB*) that drive higher levels of *galK* expression across diverse *Vibrio* species, enabling more reliable counterselection. In parallel, we generated a streptomycin-resistant *rpsL^R^*derivative of *V. diazotrophicus* to facilitate routine culturing and reduce contamination in separate experiments. This provided an opportunity to incorporate the well-established *rpsL*-mediated counterselection strategy into our toolkit. Because the *rpsL* promoter exhibits conservation comparable to that of *rpoD*, the system was broadly compatible with all streptomycin-resistant *Vibrio* species tested, eliminating the need to customize each backbone with the native *rpsL*^S^ allele. To further minimize background colony formation, we designed both pVDOG and pVRPSL to replace the target locus with a chloramphenicol resistance cassette, thereby selecting against wild-type revertants that might otherwise persist through counterselection. Following successful allelic exchange, this antibiotic cassette was removed using Flp recombinase expressed from the *Vibrio*-adapted, temperature-sensitive pVCM-*flp* vector, generating near-seamless mutants suitable for subsequent rounds of genetic manipulation.

Additionally, we developed a set of mobilizable plasmids that enable rapid tagging of *Vibrio* species with fast-maturing fluorescent proteins for straightforward visualization. These plasmids (pVFHR-4/5) are readily transferred and stably maintained in both *pir*-derived *E. coli* strains by the *oriV*_R6Kγ_, and in a broad range of *Vibrio* species via the pES213 derived high copy number origin (*oriV_Vibrio_*). Depending on the target species, these plasmids can be introduced by either conjugation or electroporation.

These counterselection strategies and tagging methods have broad applicability across the genus. Numerous *Vibrio* spp. that are not human pathogens nevertheless cause substantial economic losses in aquaculture (3, 10), contribute to coral reef decline (74, 75), and are emerging as model systems for marine microbiology (76, 77). Despite these impacts, the molecular bases of pathogenesis and host association in many understudied species remain poorly defined. The platform described here is designed to be tunable across species with divergent promoter architectures. All elements in the pVDOG, pVRPSL, and pVFHR derivatives can be excised individually or in combination using an optimized MCS with restriction enzymes active in rCutSmart buffer, enabling rapid reconfiguration. When minimizing hands-on time is a priority and a modest increase in background colonies is acceptable, a preferred counterselection cassette can be cloned into pSR6KO-1/-2, and homology arms can be assembled in a single step by overlap-extension PCR to achieve comparable outcomes (data not shown). In addition, leaving a selectable cassette between homology arms (e.g., *cat*) permits seamless incorporation of further genetic modules (*e.g.,* such as *cas9* expression cassettes or T7 RNA polymerase) thereby simplifying subsequent rounds of genome engineering as new techniques are adopted.

The systems described here each have specific limitations. The pVRPSL approach requires the target strain to carry an *rpsL*^R^ allele. In practice, this mutation was straightforward to induce in the species tested, with most strains acquiring resistance after one to five passages. The pVDOG system performed well, and we did not observe a high rate of inactivating mutations in *galK*; however, as reported previously, increasing DOG-2 concentrations can enhance recombination frequencies (57). One practical drawback is cost: DOG-2 is substantially more expensive than streptomycin, increasing per-plate expenses for counterselection. Tolerance can also arise in both systems, through additional *rpsL* mutations that confer complete resistance to ribosome-targeting antibiotics (61), or through the need for progressively higher DOG-2 concentrations to maintain recombination yields, so performance should be monitored carefully, particularly when constructing multi-locus knockouts in a single strain. Expression from pVFHR plasmids may vary by host background, resulting in different levels of visible fluorescence (data not shown). Endogenous regulation can further modulate output, as observed when replacing *rpoD* with *gyrB* promoters in the most expressive pVFHR backbones; in such cases, promoter swapping increased expression and should be considered during plasmid selection.

In summary, the tools and strategies presented here substantially expand the genetic toolkit for the *Vibrionaceae* by enabling efficient, flexible, and broadly applicable genome manipulations in both model and non-model species. By integrating robust counterselection cassettes, conserved promoters, a near-scarless Flp–FRT excision system, and versatile fluorescent-tagging plasmids, we address long-standing barriers posed by halophily, nuclease-mediated DNA degradation, and unreliable counterselection. These approaches support targeted, multi-locus genome editing and rapid visualization of diverse strains, providing a platform that can be adapted readily across phylogenetically distant taxa. Although each system has limitations, their modular design allows straightforward customization to meet species-specific challenges and to incorporate future genetic technologies. Collectively, this work establishes a foundation for deeper investigation into *Vibrio* biology, host interactions, and environmental adaptation, and equips the research community with powerful methods to interrogate pathogenicity, ecology, and functional diversity across this globally important bacterial lineage.

## Acknowledgements

This work is supported by an NSF award (2131297) to KMB. The authors would like to thank Dr. Mark Liles for providing the pCMT-flp plasmid.

## Supplemental Figure Legends

**Supplemental Figure 1. Mutations in the *rpsL* genes yield streptomycin resistance in five *Vibrio* species.** A. *Vibrio* strains were passed through sub-lethal concentrations of streptomycin to induce antibiotic resistance. Zones of inhibition surrounding a circular disc of Whatman paper (absorbing 10 µL of a 50 mg/mL stock solution streptomycin) were measured in wild-type (WT) and streptomocyin resistant (strep^R^) strains. All assays were performed in biological replicates. B. Streptomycin resistance is associated with mutations in key amino acids encoded by *rpsL* genes. The *rpsL* genes were amplified and sequenced from wildtype and strep^R^ *Vibrio* strains generated in (**A**) (*V. coralliilyticus* [V.cor], *V. diazotrophicus* [V.dia], *V. paucivorans* [V.pau], *V. mediterranei* [V.med] and *V. splendidus* [V.spl]). The predicted amino acid sequence alignment is shown. Non-synonymous mutations are highlighted in yellow. The two most common mutations occurred at positions K42 and R80. **C. Streptomycin resistance does not influence bacterial growth curves**. Growth curves were generated by culturing *Vibrio* strains at 25 C with orbital shaking for 16 hr. Absorbance measurements (O.D_.600_) were taken every five minutes. The value shown is the average of three independent biological replicates performed in three technical replicates. Standard deviation is indicated by the lighter colored shading surrounding each growth curve line.

**Supplemental Figure 2: A multilocus sequence analysis phylogenetic tree using eight conserved genes decreases resolution within the clade.** A maximum likelihood tree was constructed using the amino acid sequences of eight commonly used housekeeping present in single copies in all *Vibrio* species analyzed (Supplemental Table 4). This tree is consistent with that generated using the 25 longest single-copy orthologs (Figure 5). Bootstrap values shown were calculated using 1,000 replicates; only values greater than 75 are shown.

**Supplemental Figure 3: Successful recombination of the pVFI series does inhibit growth. A. Confirmation of integration of pVFI1-9**. Integration PCRs using a genomic flanking primer and an internal plasmid primer (Supplemental Table 3) are shown for all *V. diazotrophicus* strains harboring pVFI1–9. These strains were subsequently used to assess promoter-driven *gfp* transcript levels. A final lane contains a wild-type control amplified with the same primer set. **B. Integration does not affect growth rate.** Integrant strains and the wild-type control were cultured to stationary phase in parallel, and optical density at 600 nm (OD₆₀₀) was monitored overnight using a microplate reader. Each curve represents the mean of three biological replicates, with each biological replicate measured in technical triplicate.

## Supplemental Tables

*Supplemental Table 1: Strains used in this study*

*Supplemental Table 2: Primers used in this study*

*Supplemental Table 3: Plasmids used in this study*

*Supplemental Table 4: Bacterial genes used in phylogenetic analysis*

